# B-1a cells acquire their unique characteristics by bypassing the pre-BCR selection stage

**DOI:** 10.1101/214908

**Authors:** Jason B Wong, Susannah L Hewitt, Lynn M Heltemes-Harris, Malay Mandal, Kristen Johnson, Klaus Rajewsky, Sergei B Koralov, Marcus R Clark, Michael A Farrar, Jane A Skok

## Abstract

B-1a cells are long-lived, self-renewing innate like B cells that predominantly inhabit the peritoneal and pleural cavities. In contrast to conventional B-2 cells they have a receptor repertoire that is biased towards bacterial and self-antigens, promoting a rapid response to infection and clearing of apoptotic cells. Although B-1a cells are known to primarily originate from fetal tissues the mechanisms by which they arise has been a topic of debate for many years. Here we show that in the fetal liver (FL) versus bone marrow (BM) environment, reduced IL-7R/STAT5 levels promote immunoglobulin kappa (*Igk*) recombination at the early pro-B cell stage. As a result, B cells can directly generate a mature B cell receptor (BCR) and bypass the requirement for a pre-BCR and pairing with surrogate light chain (SLC). This ‘alternate pathway’ of development enables the production of B cells with self reactive, skewed specificity receptors that are peculiar to the B-1a compartment. Together our findings connect seemingly opposing models of B-1a cell development and explain how these cells acquire their unique properties.

## INTRODUCTION

B-CLL is the most common form of adult leukemia in the western world^1^. In the early 1980s, the T-cell antigen CD5 (Ly-1) was identified on the surface of cancerous B-cells in patients with B-CLL^2,3^. This observation led to the search for normal CD5^+^ B-cell counterparts to potentially determine the cancer cell of origin. As a result of these efforts, CD5^+^ B cells, otherwise known as B-1a cells, were discovered in mice ^4,5^. Further characterization of CD5^+^ B-1a B cells revealed that they are long-lived, self-renewing cells that predominantly reside in the pleural and peritoneal cavities where they produce natural polyreactive IgM antibodies with a biased, autoreactive repertoire. In contrast to conventional B-2 cells, B-1a cells produce antibodies with reduced junctional diversity and less somatic hypermutation ^6^. Furthermore, *Igh* V_H_ gene rearrangements favor V_H_12 segment usage ^7^, generating antibodies that interact with phosphatidylcholine (PtC), a major lipid in the protective mucus layer of the gastrointestinal tract that is also present in the membranes of diverse bacteria. Thus, the B-1a receptor repertoire is biased towards bacterial and self-antigens, which is important for mounting a rapid immune response to infection and in the clearing of apoptotic cells^8–10^. Because B-1a cells are found in preimmune mice, they function as an important first line of defense against bacterial pathogens. These characteristics distinguish B-1a cells from conventional B-2 cells, which have a highly diverse receptor repertoire that is important for mediating adaptive immunity.

Although B-1a cells were discovered in the early 1990s, their origin has been hotly debated since, and despite the efforts of numerous labs this remains an unresolved issue. The controversy has mainly been centered on two opposing models, the lineage model and the selection model. The lineage model proposes that a distinct B-1 progenitor cell gives rise to B1a cells, while the selection model favors the idea that a common B cell progenitor can acquire a B-1a or a B-2 fate depending on the type of antigen it recognizes ^9,11^. Support for the lineage model comes from early reconstitution experiments, which reveal that fetal tissues are much more efficient at generating B-1a cells in irradiated recipient mice than adult bone marrow counterparts^12^. Furthermore, the first wave of B-1a cells was shown to originate in early embryos in an HSC-independent manner ^13–17^. However, cellular barcoding experiments demonstrate that a single progenitor cell can give rise to both B-1a and B-2 cells ^18^ challenging the notion that B-1a cells arise from a distinct lineage. Moreover, the finding that B-1a cells have a restricted and biased receptor repertoire provides support for a selection model ^9,19^.

Investigations have also focused on the molecular switches that influence CD5^+^ B-1a cell fate. In this context, a lin28b/let7 axis has been characterized ^20–22^ which impacts B-1a development. The lin28b and let7 miRNAs respectively promote and inhibit the expression of the transcription factor, *Arid3*a, which in turn can drive the development of CD5^+^B-1a cells ^21,22^. Nonetheless, the B cell receptor (BCR) repertoire of the resulting cells do not include PtC specific antibodies that are characteristic of a typical B-1a cell compartment ^21^. Thus, enforced expression of *Arid3a* fails to fully explain how B-1a cells develop. Another transcription factor, BHLHE41 has also been shown to be important in B-1a cell biology ^23^. Specifically, cells deficient in this transcription factor lose B-1a cells expressing V_H_12/V_k_4 PtC specific receptors, have impaired BCR signaling, increased proliferation and apoptosis. BHLHE41 therefore plays an important role in B-1a maintenance by regulating self-renewal and BCR repertoire; however, it is not known whether its forced expression can drive development of these cells.

In the fetus, B cell development takes place in the liver and moves to the bone marrow after birth. Each stage of development is marked by a particular rearrangement event that drives differentiation forward. These recombination events occur in a stage specific manner. The first step involves the joining of *the immunoglobulin heavy-chain* (*Igh*) D_H_ and J_H_ gene segments within pre-pro B cells. Rearrangement continues at the pro-B cell stage where V_H_-to-DJ_H_ joining is both initiated and completed. Rearrangement of the immunoglobulin *light-chain* loci, *Igk* or *Igl* occurs at the subsequent pre-B cell stage of development. *Igh* and *Igk* rearrangement is separated by a proliferative burst of large pre-B cells that allows individual cells that have successfully rearranged their heavy-chain to clonally expand. At the following small pre-B cell stage, each B cell undergoes a distinct *Igk* recombination event ^24^. Ultimately, this results in unique heavy and light-chain pairs that expand the antigen receptor repertoire. The successful pairing of immunoglobulin heavy-chain with surrogate light chain (SLC) forms the pre B cell receptor (pre-BCR), which is required for expansion of large pre-B cells and subsequent differentiation to the small pre-B cell stage where *Igk* recombination occurs. Since SLC pairs poorly with autoreactive heavy-chains, the pre-BCR provides a mechanism for negative selection of self-reactive B cells ^25,26^.

However as noted early on, in a small fraction of B cell progenitors in the bone marrow *Igk* rearrangements occur prior to rearrangements at *Igh* and independent of SLCs^27–30^. In addition, in the absence of SLCs and thus pre-BCR expression, an autoreactive BCR can drive B cell development efficiently to the stage of the immature B cell, where BCR diversification and counterselection of autoreactivity is achieved through the process of receptor editing. This has led to a model in which early B cell development is driven by a positive signal from the pre-BCR (in the majority of progenitors) or an autoreactive BCR (in a minority of cells), with the pre-BCR having evolved as a surrogate autoreactive BCR ^31,32^.

Downstream of pre-BCR signaling, IL-7 receptor (IL-7R) signaling is extinguished at the small pre-B cell stage. This is noteworthy because the IL-7R signaling pathway is responsible for directing the sequential ordering of recombination events, i.e. on *heavy-chain gene* followed by *light-chain gene* rearrangement in classical B cell development ^33,34^. IL-7 is a cytokine secreted by stromal cells within the bone marrow where development takes place throughout the post-natal and adult life of the animal. In murine bone marrow, development past the pre-pro-B cell stage stringently requires IL-7R ^35,36^. In contrast, B cell development within the fetal liver can occur independent of IL-7 ^37,38^. In terms of directing recombination, IL-7 and its downstream signaling component STAT5, have been shown to promote *Igh* accessibility and recombination ^39–41^ while actively inhibiting recombination of the *Igk* locus ^33,42^. Activated STAT5 enters the nucleus and forms a complex with PRC2/EZH2 which binds to the intronic enhancer of *Igk* (iEκ) and induces H3K27me3-mediated repression to inhibit recombination of this locus ^43,44^. Strikingly, conditional deletion of STAT5 within pro-B cells results in increased *Igk* recombination at the pro-B cell stage ^34^.

It is known that B-1a cells originate primarily from fetal tissues, however it remains unclear what pathways drive B-1a cell development and lead to the acquisition of their unique characteristics. In addition, it has been shown that B-1a cells are efficiently generated in *IL-7^-/-^* and *IL-7R^-/-^* mice albeit at reduced numbers compared to wild-type ^37,38,45,46^. Given our finding that fetal liver (FL) pro-B cells have less active cytoplasmic pSTAT5 than bone marrow (BM) pro-B cells ^47^, we asked whether this could influence V(D)J recombination and B cell development. Here we show that low levels of IL-7R/pSTAT5 signaling in the fetal liver environment promote an ‘alternate pathway’ of B cell development in which increased *Igk* rearrangement occurs at the pro-B cell stage of development. Productive rearrangement of *Igh* and *Igk* at this stage leads directly to cell surface expression of a mature BCR, bypassing the requirement for SLC and selecting autoreactive receptors that are characteristic of B-1a cells. Indeed, extending earlier work^27^ we demonstrate that while SLC-independent development leads to a significant reduction in B-2 cell numbers, B-1a cells with their characteristic V_H_12 anti-phosphatidylcholine bias are still efficiently generated. Thus, reduced IL-7R/STAT5 signaling promotes an alternate pathway of development which favors the production of B-1a cells with self-reactive receptors characteristic of this B cell subset. Together these data connect opposing models of B-1a cell development and explain how these cells acquire their unique properties.

## RESULTS

### Early *Igk* recombination is increased in fetal liver versus bone marrow pro-B cells as a result of reduced STAT5-mediated repression

Fetal liver (FL) and bone marrow (BM) derived progenitor cells are differentially dependent on IL-7 for development and we have found that fetal liver derived pro-B cells have significantly lower levels of the downstream signaling component, phospho-STAT5 compared to their bone marrow derived pro-B cell counterparts ^47^. Thus, we asked whether alterations in phospho-STAT5 levels correlate with altered regulation of *Igk* recombination in these two anatomic locations. To explore this, we analyzed recombination from *ex vivo* derived bone marrow (5-6wk adults) and fetal liver (E17-18) pro-B cells using semi-quantitative PCR analysis (**Fig. 1a**). This assay uses a degenerate Vκ primer with a common reverse primer downstream of Jκ gene segments to measure recombination of a Vκ with each of the four Jκ segments^48^. As shown in Figure. 1a, FL pro-B cells have increased *Igk* recombination relative to BM pro-B cells.

**Figure 1.**
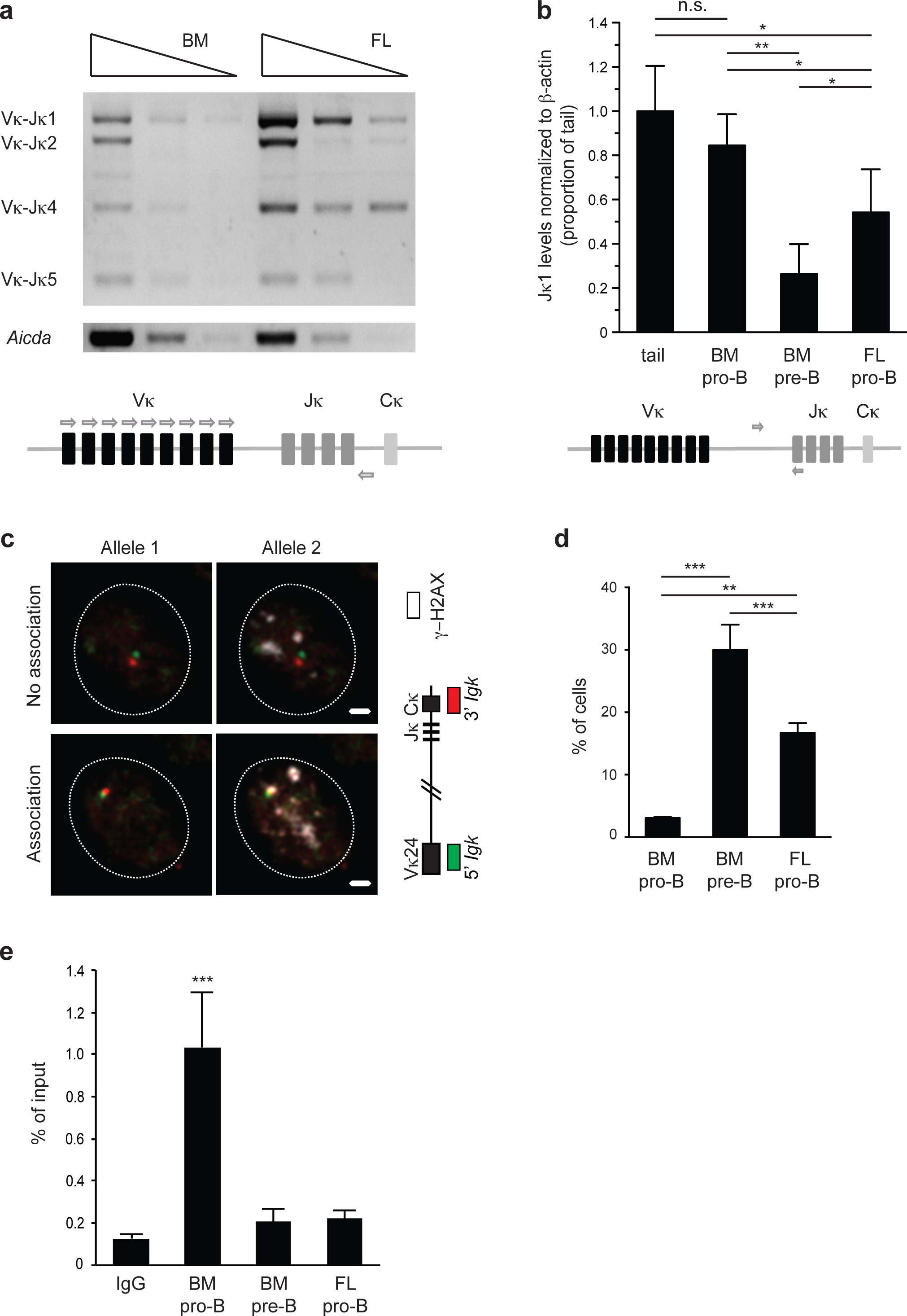
Early *Igk* recombination is increased in fetal liver versus bone marrow pro-B cells as a result of reduced STAT5-mediated repression. (**a**) Semi-quantitative PCR performed on ex vivo derived BM (6 weeks) and FL pro-B (E17.5) to assess rearrangement of Vκ to each functional Jκ. Each lane represents 3-fold serial dilutions of input DNA. DNA levels were normalized to *Aicda* levels, which is located on the same chromosome as *Igk*. Cartoon outlining the location of the primers on the *Igk* locus (**b**) Recombination with Jκ1 was quantified by qPCR using primers specific for the unrearranged germline sequence. DNA is quantified as a ratio between the single copy *β*-*actin* gene and the *Ig*κ germline product and shown as a proportion of tail DNA. Cartoon outlining the location of the primers on the *Igk* locus. Data is displayed as an average of two independent experiments and error bars represent the standard deviation. P-values were calculated using two-tailed T-tests (**c**) 3-D immuno-DNA FISH was performed on *ex vivo* sorted B cells using BAC probes specific to the distal Vκ24 (RP23-101G13) gene region and the Cκ region (RP24-387E13), shown in red and green respectively, in conjunction with an antibody to the phosphorylated form of γ-H2AX in white. Representative images of a B cell with no γ-H2AX associated with *Igk* alleles (top) and with γ-H2AX associated with one *Igk* allele (bottom). Scale = 1μm. (**d**) Percentage of B cells (BM pro-B, BM pre-B and FL pro-B) with at least one *Igk* allele associated with a γ-H2AX focus. P-values were calculated using two-tailed Fisher's exact tests. (**e**) STAT5-ChIP qPCR of iEκ on *ex vivo* sorted cells. Data is displayed as an average of 3 biological replicates as a proportion of input DNA and error bars represent the standard deviation. P-values were calculated using two-tailed T-tests. For all P-values, *significant (0.05–0.01) **very significant (0.01-0.001), ***highly significant (<0.001).

To address the question of altered *Igk* recombination between these two cell types in a more quantitative manner, we made use of a real-time PCR assay that quantities Jκ1 rearrangement by assessing the retention of germline *Igk* ^49^. In this assay, a product cannot be generated after recombination because the sequence to which the upstream primer binds is lost or inverted during the recombination process. Quantitation of the amplified product is determined as a ratio between the single copy gene *β*-*actin* and the germline *Igk* band. Control tail DNA, in which no *Igk* rearrangement occurs, is set at 100%. Consistent with the semi-quantitative experiments described above, fetal liver pro-B cells have lower levels of *Igk* germline retention (61%) relative to bone-marrow pro-B cells (87%) (**Fig. 1b**).

To determine the proportion of cells that are actively undergoing recombination at a single cell level, we used three-dimensional (3-D) immuno-flourescent *in situ* hybridization (FISH) assay that visualizes recombination by the association of γ-H2AX foci with the individual alleles of antigen receptor loci. γ-H2AX DNA repair foci have been shown to be associated with antigen receptor loci undergoing recombination ^50,51^. In these experiments, we used two *Igk* BAC DNA probes that hybridize to the distal Vκ24 (RP23-101G13) gene region and the Cκ region (RP24-387E13) in combination with an antibody against the phosphorylated form of γ-H2AX (**Fig. 1c**). We found significantly increased association of γ-H2AX foci with the *Igk* locus in fetal liver versus bone marrow derived pro-B cells (16.7% versus 3.0%). Consistent with our previous experiments the frequency of DNA breaks in FL pro-B cells (16.7%) is again at an intermediate level compared to BM pre-B cells (30%) where *Igk* recombination typically occurs (**Fig. 1d**). These numbers reflect the combined data from two independent experiments (**Supplementary Table S1**).

Antigen receptor loci undergo large-scale contraction through chromatin looping which enables rearrangement between widely dispersed gene segments. Locus contraction is tightly linked both to recombination status and usage of distal gene segments ^48,52^. To address the contraction status of the *Ig* locus we measured the distance separating the distal Vκ24 and Cκ probes in interphase cells. Our analyses indicate that FL pro B cells are significantly more contracted than wild-type (WT) double positive (DP) T cells, a cell type in which *Ig* rearrangement does not occur (**Supplementary Fig. 1a,b**). Indeed the level of contraction is comparable to that seen in wild-type bone marrow derived pre-B cells where *light-chain* rearrangement is known to occur.

It is known that activated STAT5 forms a complex with PRC2 and binds to the intronic enhancer of *Igk*, iEκ to inhibit *Igk* recombination^43,47,53^. To determine if the increase in early *Igk* rearrangement could be caused by reduced STAT5-mediated repression, we assessed how much STAT5 is bound to *Igk* via STAT5-ChIP qPCR. As shown in **Fig. 1e**, we found that FL pro-B cells have significantly less STAT5 bound to iEκ compared to BM pro-B cells and binding more closely resemble levels found in BM pre-B cells where *Igk* recombination normally occurs. Taken together, these findings indicate that in the FL versus BM environment *Igk* rearrangement occurs at higher levels in pro-B cells due to a decrease in STAT5-mediated repression.

### Pro-B cells that undergo early *Igk* rearrangement can bypass the pre-BCR checkpoint and make B cells that express a mature BCR

Classical bone marrow B cell development starts with *Igh* recombination at the pre-pro to pro-B cell stages followed by *Igk/Igl* recombination at the small pre-B cell stage. However, JHT mice that lack J_H_ segments as well as the *Igh* intronic enhancer, and are therefore unable to undergo *Igh* recombination, can rearrange *Igk/Igl* independent of *Igh* rearrangement (**Supplementary Fig. 2**)^28,54,55^ This is also true for SLC deficient mice and a fraction of B cells undergoing development in the bone marrow of wild-type mice ^56^. Given our finding that a significant number of pro-B cells rearrange *Igk* in the fetal liver environment, we reasoned that productive *heavy* and *light-chain gene* rearrangement in pro-B cells could lead to expression of a mature BCR, bypassing the requirement for a pre-BCR. To address this question as a proof of principle, we examined fetal liver B cells from mice that are deficient for a component of surrogate light chain, Lambda5 (*Igll1^-/-^*), in conjunction with a transgene that has a prearranged functional heavy-chain (B1.8). Whereas *Igll1^-/-^* mice display a strong developmental block at the pro-B cell stage and made very few IgM^+^ cells (0.44% of B cells). On the other hand, B1.8 and *Igll1^-/-^*; B1.8 mice produce similarly high proportions of IgM+ B cells (12 and 13 % respectively), which are also IgK^+^. Importantly, B1.8 mice have a substantial pre-B cell compartment which is not detected in *Igll1^-/-^*, B1.8 mice despite similar IgM^+^ B cell output (**Fig. 2**). This shows that having a productive heavy and light-chain can bypass the requirement for pairing with SLC to form immature B cells that express the mature BCR. Together this data shows that in the fetal liver, expression of *Igh* and *Igk* at the pro-B cell stage can bypass the developmental block induced by surrogate light chain deficiency by forming B cells with a mature BCR.

**Figure 2.**
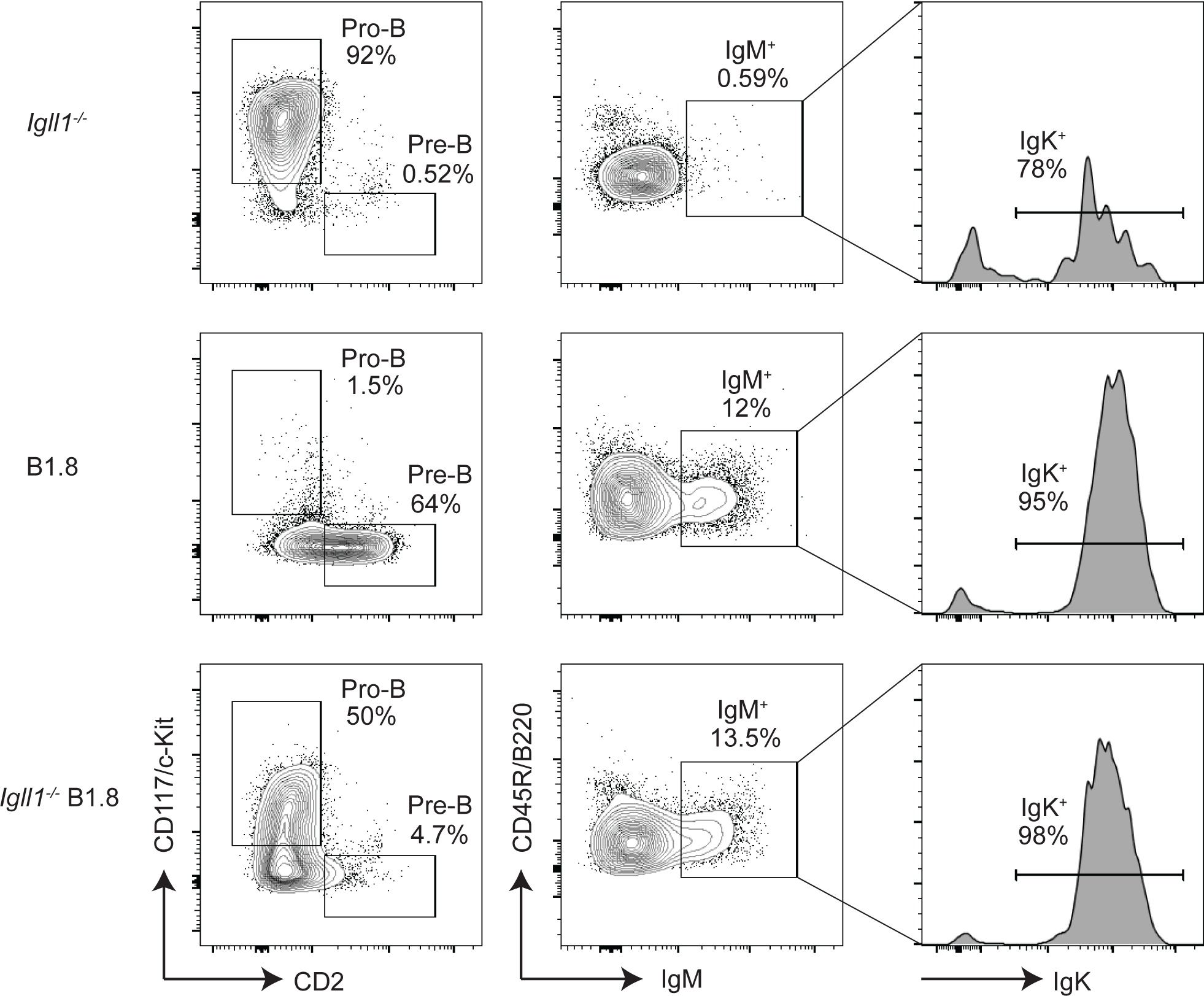
Pro-B cells that undergo early *Igk* rearrangement can bypass the pre-BCR checkpoint and make B cells that express a mature BCR. (**a**) Representative flow cytometry plots of fetal liver E17.5 cells from *Igll1^-/-^* and *Igll1^-/-^*; B1.8 littermates. Pro-B and pre-B cell gates are displayed as a percentage of CD19^+^, IgM^-^ cells (left), IgM^+^ are displayed as a percentage of CD19^+^ cells (middle), and IgK^+^ histograms are displayed as a percentage of CD19^+^, IgM^+^ cells. (**b**) Graphical representation of IgM^+^ cells as a percent of CD19^+^ B cells. Data are displayed as the average of 3 independent experiments and error bars show the standard deviation. P-values were calculated using a two-tailed T-test, *significant (0.05-0.01) **very significant (0.01-0.001), ***highly significant (<0.001).

### B-1a cells are efficiently generated in surrogate light deficient mice

B-1a cells are primarily generated from fetal tissues, and we have shown that FL derived B cells frequently undergo early *Igk* recombination allowing cells to bypass the pre-BCR selection stage. Thus we next asked whether an absence of surrogate light chain could impact the generation of B-1a cells ^12,57^. Here we analyzed the B-2 as well as the B-1 compartment, which consists of both B-1a and B-1b cells. B-1b cells are distinct from B-1a cells in that they lack CD5 expression and have a memory function in protecting against bacterial infections ^58^. As expected, peritoneal B-2 cells were significantly decreased compared to controls. Furthermore although B-1b cells were also significantly decreased, the B-1a cell compartment remained intact. This was reflected in cell percentage as well as total cell numbers (**Fig. 3a,b**). These data indicate that while B2 and B1b cells development is strongly promoted by SLC, B-1a cells do not depend on a pre-BCR checkpoint. It should be noted that B1.8 transgenic mice could not be used to analyze the B-1a cell compartment in surrogate light chain deficient mice because the B1.8 pre-rearranged heavy-chain does not support B-1a cell development ^59^.

**Figure 3.**
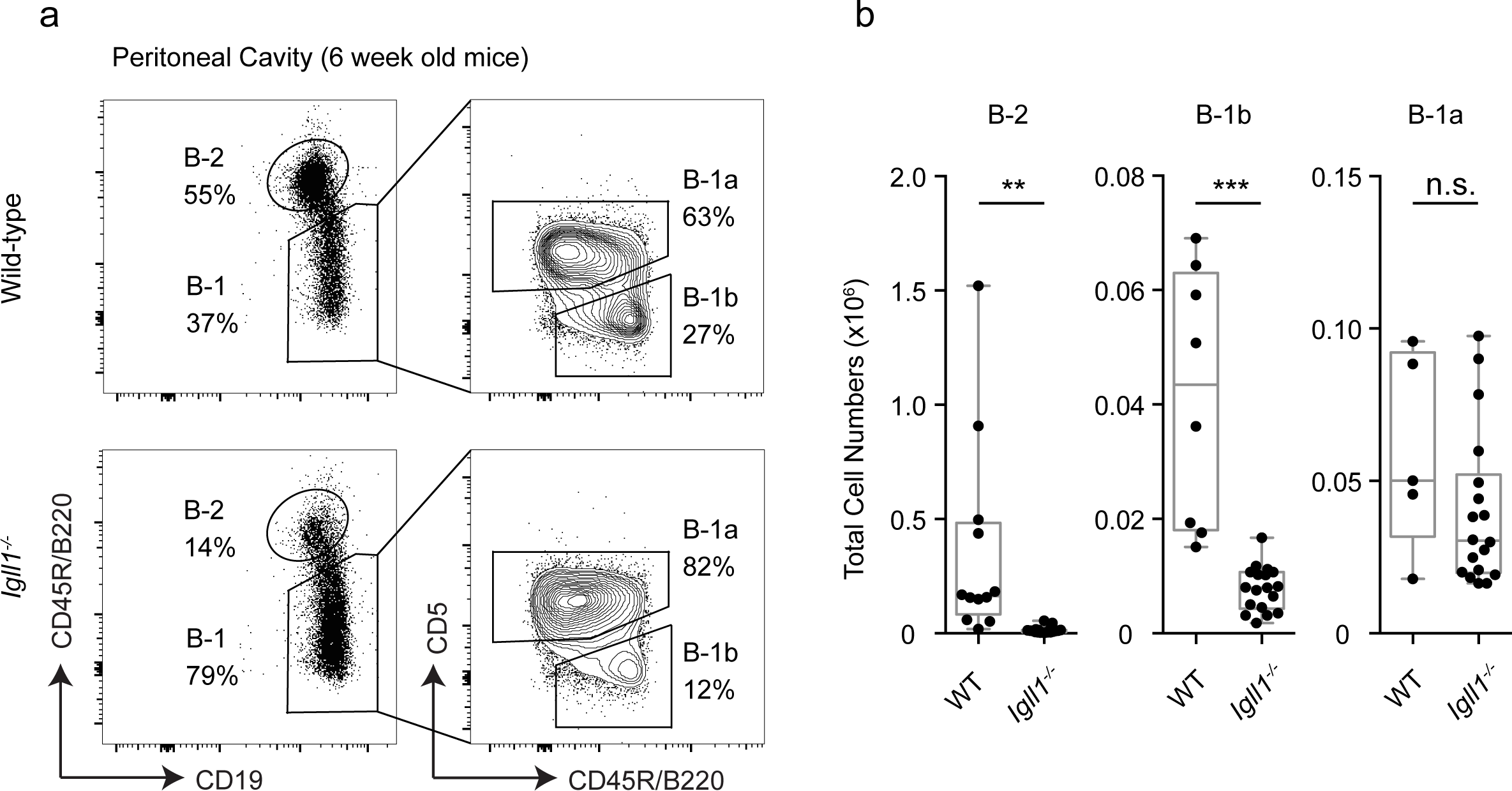
Pro-B cells that undergo early *Igk* rearrangement can bypass the pre-BCR checkpoint and make B cells that express a mature BCR. (**a**) Representative flow cytometry plots of peritoneal cavity cells from wild-type and *Igll1*^-/-^ mice. B-2 and B-1 cell gates are displayed as a percentage of CD19^+^ cells (left). B-1a and B-1b cell gates are displayed as a percentage of B-1 cells (right). (**b**) Total cell numbers of B-2, B-1b and B-1a cells were calculated from the peritoneal cavity of these mice. Data is displayed in box plots and each dot represents an individual mouse. P-values were calculated using two-tailed T-tests, *significant (0.05-0.01) **very significant (0.01-0.001), ***highly significant (<0.001).

Two different subsets of B-1a cells that segregate different functions of the B-1a cell compartment, have been identified by plasma cell alloantigen 1 (PC1) ^60,61^. In order to determine whether or not both of these subsets of B-1a cells were represented in B-1a cells generated in surrogate light chain deficient mice, we looked at PC1 expression. B-1a cells from *Igll1*^-/-^ mice were found to have similar proportions of PC1^lo^ and PC1^hi^ B-1a cells as in wild-type mice (**Supplementary Fig. 3**). This suggests that not only is the B-1a compartment size normal in surrogate light chain knockout mice, but PC1^lo^ and PC1^hi^ subsets are maintained.

### Constitutive phospho-STAT5 signaling selectively inhibits B-1a cell development

Our data suggest that low phospho-STAT5 signaling induces early *Igk* recombination in FL pro-B cells, which promotes the efficient generation of B-1a cells in surrogate light chain deficient mice. To further validate the role of IL-7R/STAT5 signaling in B-1a cell development, we next asked whether increased STAT5 signaling could have the opposite effect on B-1a cell development and impair the generation of this compartment. To address this question we used mice expressing a constitutive active form of STAT5 (*Stat5b-CA*) ^62^.

In these mice we found that although B-2 and B-1b cell development were not significantly affected, all three independent littermate controlled *Stat5b-CA* mice exhibited a decrease in the B-1a cell compartment. This is reflected in cell proportions as well as total cell numbers (**Fig. 4a,b**). Thus, STAT5 signaling inhibits B-1a cell development underscoring the importance of reduced IL-7R/STAT5 signaling in driving the development of these cells.

**Figure 4.**
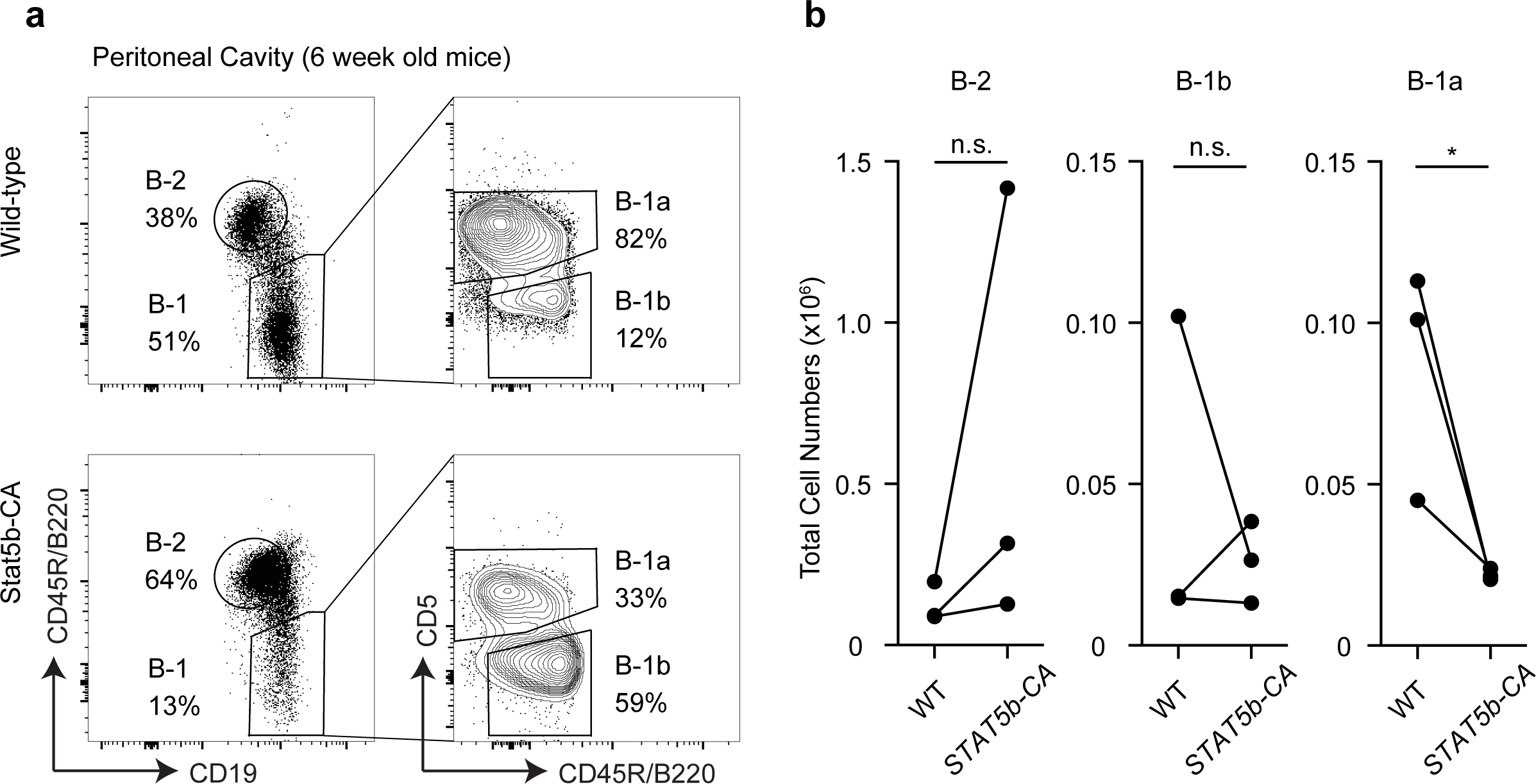
Constitutive phospho-STAT5 signaling selectively inhibits B-1a cell development. (**a**) Representative flow cytometry plots of peritoneal cavity cells from wild-type and *STAT5b-CA* mice. B-2 and B-1 cell gates are displayed as a percentage of CD19^+^ cells (left). B-1a and B-1b cell gates are displayed as a percentage of the B-1 cells (right). (**b**) Total cell numbers of B-2, B-1b, and B-1a cells were calculated from the peritoneal cavity of these mice. Each dot represents an individual mouse and lines connect pairs of littermates. P-values were calculated by paired two-tailed T-tests, *significant (0.05-0.01) **very significant (0.01-0.001), ***highly significant (<0.001).

### SLC independent B-1a cells express receptors with phosphatidylcholine (PtC) specific V_H_12 gene rearrangements

B-1a cells are known to have a strong bias for immunoglobulin rearrangements that confer specificity to phosphatidylcholine (PtC) (V_H_12/V_k_4). In addition, the pre-BCR is known to select against B-1a specific autoreactive heavy-chains as a result of a defect in pairing of the latter with surrogate light chain ^25,26^. Given these facts we next asked whether B-1a cells that develop independent of SLC could lead to the generation of B-1a specific rearrangements. To address this question we analyzed peritoneal cavity B cells for V_H_12 usage and specificity against PtC. As shown in **Fig. 5a** B-1a cells from *Igll1^-/-^* mice express V_H_12 rearranged receptors. Importantly, these V_H_12^+^ B-1a cells bind PtC containing liposomes inferring that they have the canonical V_H_12/V_k_4 gene segment bias (**Fig. 5a**). Additionally, there is not a significant difference in the proportion of B-1a cells that have specificity against PtC (**Fig. 5b**). Thus, surrogate light chain independent B-1a cell development supports the generation of these B-1a cells.

**Figure 5.**
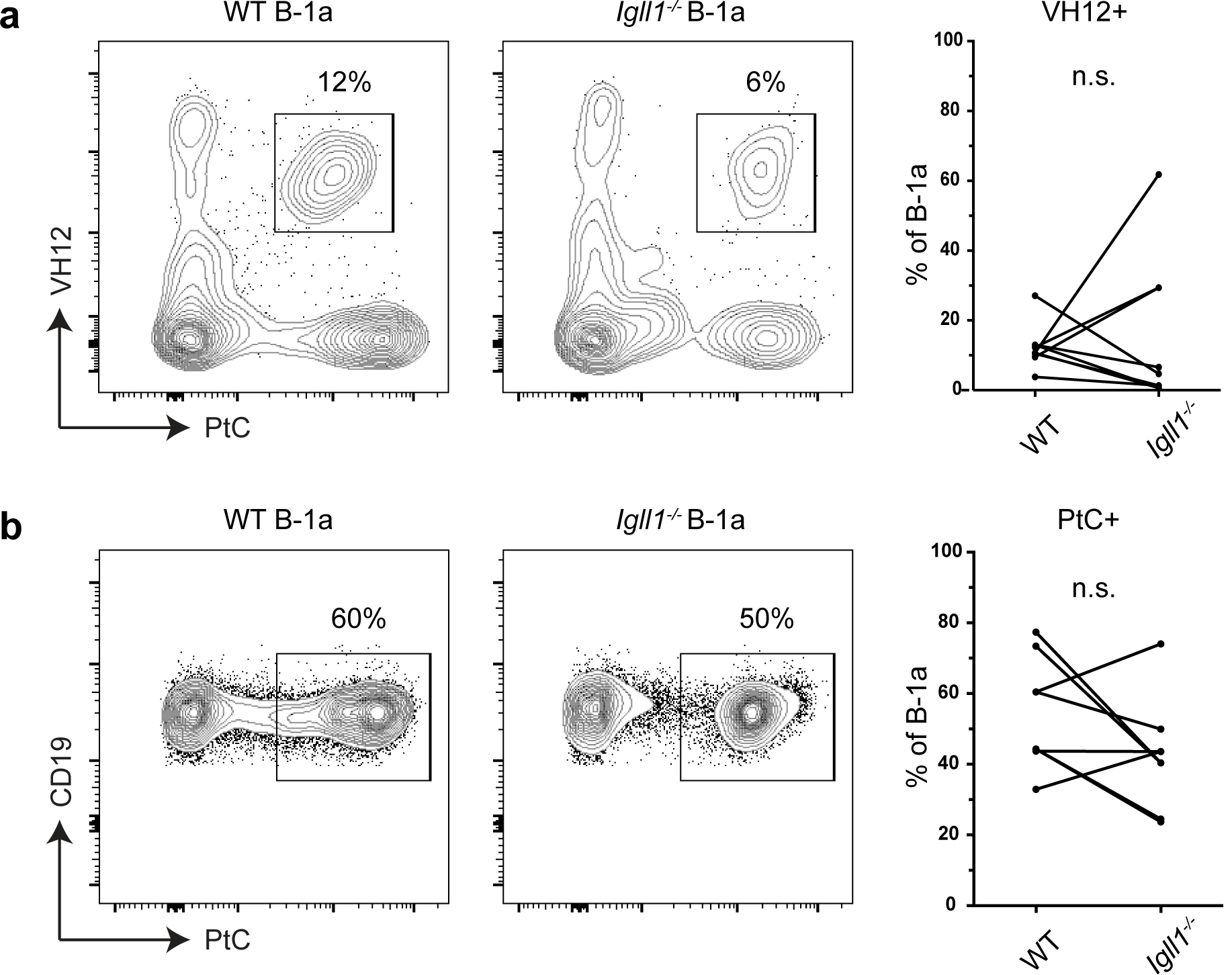
SLC independent B-1a cells express receptors with phosphatidylcholine (PtC) specific V_H_12 gene rearrangements. Peritoneal cavity B-1a cells from wild-type and *Igll1^-/-^* mice are shown that highlight the proportion of B-1a cells that are either: (**a**) VH12^+^, PtC^+^ or (**b**) PtC^+^. The left side shows representative flow cytometry plots. The right side is a graphical summary of the mice from all of the experiments. Each dot represents an individual mouse and lines connect pairs of littermates. P-values were calculated by paired two-tailed T-tests, *significant (0.05–0.01) **very significant (0.01-0.001), ***highly significant (<0.001).

### B-1a cells have a similar gene segment usage to wild-type controls

To determine how SLC deficiency affects other V_H_ gene segment usage, we sequenced the BCR repertoires of B-1a cells from WT and SLC-deficient mice. Briefly, peritoneal cavity B-1a cells were sorted, lysed and the rearranged *Igh* products amplified by PCR. Illumina adapters and barcodes were added and the rearrangements sequenced by next-generation sequencing. BCR repertoires were analyzed by IMGT/High-VQUEST and visualized using IMGT/StatClonotype^63–66^. These analyses revealed that B-1a cells from *Igll1^-/-^* mice have a similar overall V_H_ usage compared to those of wild-type mice (**Fig. 6**). This shows that B-1a cells that develop in the absence of SLC have a wide distribution of gene segment usage rather than representing the progeny of a few rare progenitors. As expected V_H_12 (V_H_12-3) usage, was found to be high in wild-type B-1a cells and levels were matched in the cells from *Igll1^-/-^* mice (**Fig. 6**) ^7^. The rearrangement studies support the data from our FACs analysis and underscore the finding that B-1a cells generated in the absence of SLC maintain a V_H_12 gene segment bias.

**Figure 6.**
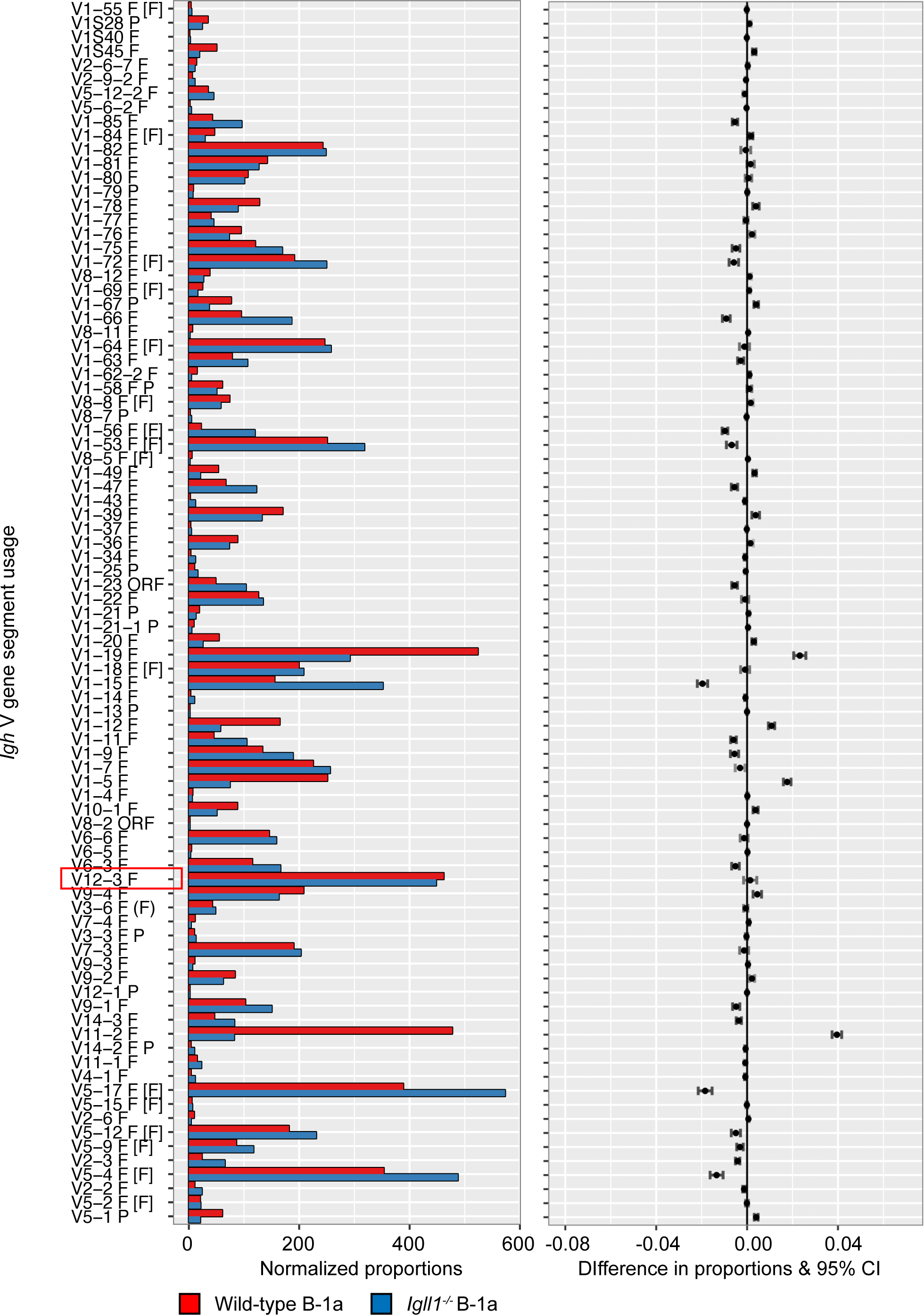
*Igll1^-/-^* B-1a cells have a similar gene segment usage to wild-type controls. Representative synthesis graphs generated by IMGT/StatClonotype that compare differences in V gene segment usage between wild-type (Red) and *Igll1^-/-^* (Blue) derived B-1a cells. The red box highlights V_H_12 (VH12-3) gene segment. This graph combines a bar graph for the normalized proportions of each gene segment and the differences in proportions with significance and confidence intervals (CI).

## DISCUSSION

Although it was known that B-1a cells predominantly develop in the fetal liver, the pathways driving development and the mechanisms underlying the generation of B cells with a repertoire skewed towards autoreactivity was not previously known. Indeed, this has been a subject of debate for many years. Here we now reveal that a reduction in IL-7 signaling in the fetal liver environment alleviates *Igk* repression and promotes early rearrangement of this locus in pro-B cells. Productive *heavy* and *light-chain* gene rearrangement at the pro-B cell stage can lead directly to the expression of a mature BCR, thereby bypassing the requirement for a pre-BCR checkpoint. Since PtC specific V_H_ gene rearrangements pair poorly with SLC ^25^ and the pre-BCR selects against autoreactive receptors ^26^, SLC independent development provides an explanation for how B cells with autoreactive, PtC specific receptors are generated. In more general terms and along Jerne’s idea that the immune system selects antibody mutants from an initial self-reactive repertoire ^67^, B cell progenitors can be initially positively selected by the expression of autoreactive BCRs or their evolutionary surrogate, the pre-BCR ^31,32^. This is followed by negative selection of autoreactivity through receptor editing. The skewing of the B-1a receptor repertoire towards autoreactivity likely reflects the imprint of the initial positive selection of B-1a cells through autoreactive BCRs rather than SLC.

The idea that B-1 cells can develop in a SLC independent manner is not novel. In fact, Kitamura *et al*. (1992) observed that in contrast to B-2 cells B-1 cells were essentially unaffected by SLC deficiency, however the distinct effect on the B-1a versus B-1b cell compartment was not investigated^27^. Our analyses demonstrate that while B-2 and B-1b cell development is severely impaired in *Igll1^-/-^* mice, B-1a cells can be generated at wild-type levels. Furthermore, SLC independent B-1a development could occur through a mechanism that involves reduced IL-7/STAT5 signaling and early *Igk* rearrangement. We found that B-1a cell development is selectively impaired in *Stat5b-CA* mice and this provides additional support for the idea that SLC independent B-1a cell development is favored by a decrease in IL-7 signaling.

Taken together our findings point to a model in which B-1a cells typically develop in a SLC-independent manner. Interestingly, earlier work analyzing expression patterns of genes relevant to rearrangement in the yolk sac (YS), para-aortic splanchnopleura (P-Sp) and spleen demonstrate that at E9-11, early lymphoid progenitor cells express *Rag2* and *VpreB* but lack expression of *Igll1*^68^. This delayed expression of the SLC component, lambda5 (*Igll1*), points to the possibility that there is an early window in which B cell development can occur in the absence of SLC. In fact, this stage coincides with the stage at which the earliest B-1 progenitor cells are detected (E9 in the YS and P-Sp)^69,70^. Thus, we speculate that the reason why the early progenitors are B-1 specific is because they develop in an SLC-independent manner. The observation that SLC deficient mice efficiently generate B-1a cells with autoreactive receptors supports this model^26^. In sum our assignment of a SLC independent pathway of B cell development to B-1a cells provides a new perspective on B-1a development, reconciling old lineage and selection models (Fig. 7)

**Figure 7.**
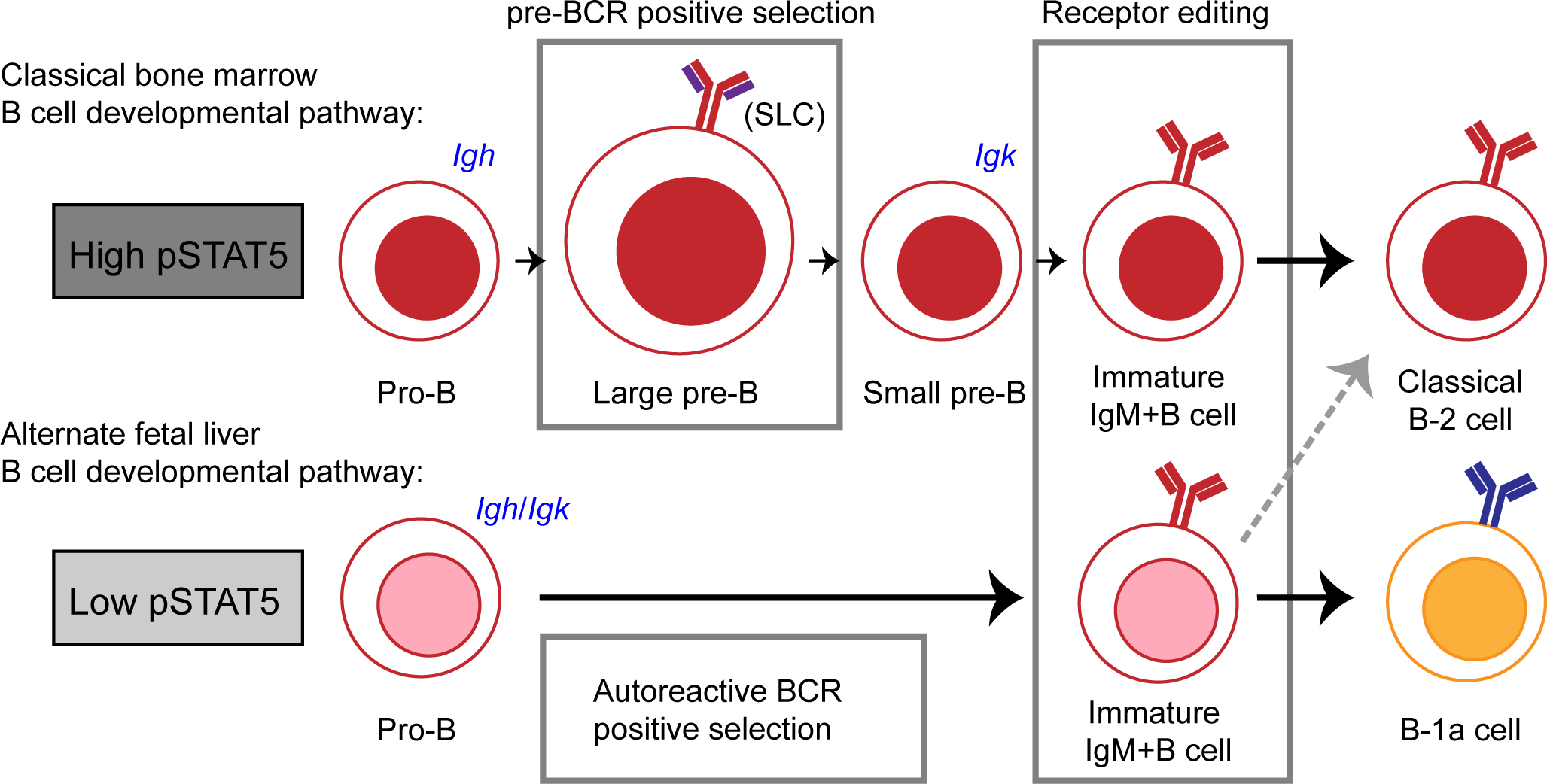
Model of classical bone marrow B cell development versus alternate fetal liver B cell development. In the classical bone marrow B cell developmental pathway, high levels of STAT5 signaling largely prevent *Igk* recombination at the pro-B cell stage. At the large pre-B cell stage, a productive heavy-chain pairs with SLC and pre-BCR signaling at this stage leads to proliferative expansion and positive selection. Autoreactive heavy-chains pair poorly with SLC and will not experience positive selection in this manner. Following *Igk* recombination at the small pre-B cell stage, self-reactive B cells can undergo receptor editing to further select against self-reactive BCRs. In the alternate fetal liver B cell developmental pathway generating B-1a cells, low levels of STAT5 signaling allow for *Igk* recombination at the pro-B cell stage, allowing the generation of B cells expressing a mature BCR instead of a pre-BCR. Here, self-reactive B cells are initially positively selected by self-antigens. Although these B cells can potentially undergo receptor editing to modify their BCRs, the initial positive selection of autoreactive B cells skews the B-1a BCR repertoire towards autoreactivity.

## Materials and Methods

### Mice

*Igll1^-/-^* mice (Jackson Laboratory 002401) were maintained on a C57BL/6 background and genotyped as previously described^27^. B1.8 mice^71^ were crossed onto a *Igll1^-/-^* background. These mice were housed and cared for in accordance with IACUC guidelines and protocols approved by NYUMC (protocol #: IA15-01468) and U of Minnesota (protocol #: 1502-32347A).

### Flow cytometry and antibodies

Bone marrow and fetal liver cell populations were isolated from C57Bl/6 mice via cell sorting and analyzed by flow cytometry. B and T cell antibodies include: anti-CD45R/B220 (RA3-6B2), anti-CD19 (1D3), anti-IgM^b^ (AF6-78), anti-IgM (II/41), anti-CD117/c-Kit (2B8), anti-CD2 (RM2-5), anti-CD25/IL2RA (PC61), anti-CD90.2/Thy1.2 (53-2.1), anti-CD5 (53-7.3), anti-VH12 (5C5)^72^ anti-TCRB (H57-597), anti-CD8a (53-6.7), anti-CD4 (RM4-5). These antibodies were obtained from either BD Pharmigen or eBioscience. Fluorescent DPOC/CHOL Liposomes were acquired from FormuMax. The gating strategy was as follows: bone marrow pro-B cells B220^+^/CD19^+^/IgM^-^/c-kit^+^/CD25^-^/IgM^-^, bone marrow pre-B cells B220^+^/CD19^+^/ IgM^-^/c-kit^-^/CD25^+^, fetal liver pro-B cells (E17.5) B220^+^/CD19^+^/IgM^-^/c-kit^+^/CD2^-^ and thymic DP cells Thy1.2^+^/TCRB-/CD19^-^/CD4^+^/CD8^+^. Fetal liver pro-B cells were sorted on CD2^-^ instead of CD25^-^ because CD25 is not expressed in fetal B cells. CD2 expression is correlated with cytoplasmic μ heavy-chain and can be used to identify pre-B cells and efficiently sort pro-B cells from the fetal liver ^73^. From the peritoneal cavity B-2, B-1a and B-1b were identified as followed: B-2 cells B220^hi^/CD19^+^, B1a cells B220^lo^/CD19^+^/CD5^+^ and B-1b cells B220^lo^/CD19^+^/CD5^-^. To obtain fetal liver precursor cells, fetal livers from E15.5-17.5 were subject to lineage depletion using the following biotin labeled antibodies: anti-CD4 (RM4-5), anti-CD8a (53-6.7), anti-TCRγθϏϏϏ, anti-CD49b (DX5), anti-Ly6G/Ly6C (RB6-8C5), anti-CD11c (N418), anti-CD11b (M1/70), anti-F4/80 (BM8), anti-Ter-119 (TER-119) followed by a negative selection using streptavidin magnetic rapidspheres (STEMCELL Technologies). The remaining cells were sorted as: Lin^-^/CD19^-^/IgM^-^/c-kit^+^/CD2-/B220^lo^. Cells were sorted using a FACSAria I (BD). Data were also collected on an LSR II (BD) and analyzed using FlowJo software.

### Semiquantitative *Igk* rearrangement and Jκ1 germline retention analysis

DNA was prepared from sorted cells by proteinase K digestion at 55C for 4 hours, 85C for 10 mins. VκJκ joints were amplified using a degenerate Vκ primer (5’-GGCTGCAGSTTCAGTGGCAGTGGRTCWGGRAC-3’), and a Jκ5 primer (5’-ATGCCACGTCAACTGATAATGAGCCCTCTCC-3’). *Acidia* gene was amplified as a loading control (AID F: 5’-GCCACCTTCGCAACAAGTCT-3’, AID R: 5’-CCGGGCACAGTCATAGCAC-3’). Bands were run through a 1.5% agarose gel electrophoresis and imaged using a ChemiDoc XR+ (Bio-Rad). Germline retention analysis of Jκ1 was carried out by real-time quantitative PCR analysis as previously described^49^.

### Immuno-DNA FISH

Combined detection of γ-H2AX and *Ig* loci was carried out on cells adhered to poly-L lysine coated coverslips as previously described^50^. Cells were fixed with 2% paraformaldehyde / PBS for 10 minutes and permeabilized for 5 minutes with 0.4% Triton / PBS on ice. After 30 minutes blocking in 2.5% BSA, 10% normal goat serum and 0.1% Tween-20 / PBS, H3S10ph staining was carried out using an antibody against phosphorylated serine-10 of H3 (Millipore) diluted at 1:400 in blocking solution for one hour at room temperature. Cells were rinsed 3 times in 0.2% BSA, 0.1% Tween-20 / PBS and incubated for one hour with goat-anti-rabbit IgG Alexa 488 or 594 or 633 (Invitrogen). After 3 rinses in 0.1% Tween-20 / PBS, cells were post fixed in 3% paraformaldehyde / PBS for 10 minutes, permeabilised in 0.7% Triton-X-100 in 0.1M HCl for 15 minutes on ice and incubated in 0.1 mg/ml RNase A for 30 minutes at 37°C. Cells were then denatured with 1.9 M HCl for 30 minutes at room temperature and rinsed with cold PBS. DNA probes were denatured for 5 minutes at 95°C, pre-annealed for 45 minutes at 37°C and applied to coverslips which were sealed onto slides with rubber cement and incubated overnight at 37°C. Cells were then rinsed 3 times 30 minutes with 2x SSC at 37°C, 2x SSC and 1x SSC at room temperature. Cells were mounted in ProLong Gold (Invitrogen) containing DAPI to counterstain total DNA.The *Ig* locus was detected using BAC DNA probes that hybridize to the distal Vκ24 gene region (RP23-101G13) and the Cκ region (RP24-387E13), in combination with an antibody against the phosphorylated form of γ-H2AX. BAC probes were directly labeled by nick translation with dUTP-A594 or dUTP-A488 (Invitrogen). FISH for locus contraction was conducted as previously published^50^.

### Confocal microscopy and analysis

Cells were analyzed by confocal microscopy on a Leica SP5 Acousto-Optical Beam Splitter system. Optical sections separated by 0.3 μm were collected. Analysis on cells included those in which signals from both alleles could be detected which encompassed 90-95% of the total cells imaged. Further analysis was carried out using ImageJ. Alleles were defined as colocalized with γ-H2AX if the signals overlapped. Sample sizes typchically included a minimum of 100 cells and experiments were repeated at least two or three times. Statistical significance was calculated by χ^2^ analysis in a pair-wise manner. For locus contraction, distances were measured between the center of mass of each BAC signal. Significant differences in distributions of empirical inter-allelic distances were determined by a nonparametric two-sample Kolmogorov-Smirnov (KS) test. To eliminate observer bias, each experiment was analyzed by at least two people.

### STAT5 ChIP-qPCR

STAT5 ChIP-qPCR was conducted as previously described^43^. Purified DNA was then analyzed by quantitative real-time PCR. Samples were analyzed in triplicate and represent three biological replicates.

### *Igh* repertoire DNA library preparation and sequencing analysis

Cells were sorted and lysed in 0.5 mg/mL proteinase K at 55C for 4 hours, 85C 10 mins. Lysed cells were used as template for HotStarTaq (Qiagen) PCR amplification. Primers used to amplify rearrangements were adapted from V_H_ FR3 and J_H_ BIOMED-2 primers^74,75^. Additional primers designed to capture some V genes missing from the BIOMED-2 study, which include V_H_12 genes (**Supplementary Table S2**). Amplified rearrangements were purified by gel extraction (Qiagen) followed by a standard end-repair reaction. Following purification steps were carried out by Ampure XP bead purification (Beckman Coulter). In order to attach Illumina-compatible adapters, samples were treated by standard dA tailing followed by adapter ligation using Quick Ligase (NEB) mixed with preannealed NEXTflex DNA Barcodes (Bioo Scientific). QC was carried out by tapestation and quantified by qPCR (Kapa Biosystems). Samples were pooled and sequenced on an Illumina HiSeq 4000 (2 x 150 PE reads). Nucleotide sequences were compared to the reference equences from IMGT, the international ImMunoGeneTics information system (http://www.imgt.org) and analyzed using IMGT/HighV-QUEST^66^, a web portal allowing the analysis of thousands of sequences on IMGT/V-QUEST^64^. IMGT/StatClonotype was used to Analyze statistically significant gene segment usage between samples^63,65^.

## ACKNOWLEDGMENTS

We would like to thank members of the Skok lab for thoughtful discussions and critical comment on the study and manuscript. We would also like to thank the NYU Flow Cytometry and Cell Sorting Center as well as the NYU Genome Technology Center, both of which are supported in part by the Cancer Center Support Grant P30CA016087 at the Laura and Isaac Perlmutter Cancer Center. JAS is supported by NIH grants R35GM122515 and R21CA188968. JBW was previously supported by the T32 CA009161 training grant (Levy) and is currently supported by the 2T32 AI100853-06 (Reizis) training grant. SLH was supported by an American Society of Hematology (ASH) Scholar Award and by a Molecular Oncology and Immunology Training Grant NIH T32. MAF is supported by NIH grants CA151845 and CA154998.

---

**Supplementary Figure 1** Fetal liver pro-B cells and bone marrow pre-B cells are similarly contracted at the *Igk* locus.(**a**) Representative confocal microscopy images showing the distance separation between the probes on *Igk* in fetal liver pro-B cells, bone marrow pre-B cells and double positive T cells. (**b**) Distances separating the two ends of the locus are displayed as a cumulative frequency curve. A left shift on the curve is indicative of closer association. Bone marrow pre-B cells (blue), fetal liver pro-B cells (green), and negative control double positive T cells (black). P-values were generated using two-sample Kolmogorov-Smirnov tests.

**Supplementary Figure 2** *Igk* recombination can occur independent of *Igh* recombination. Semi-quantitative PCR performed on ex vivo derived fetal liver pro-B cells (E17.5) from wild-type and JHT mice. Each lane represents 3-fold serial dilutions of input DNA. DNA levels were normalized to iEκ, which is also located on the same chromosome as *Igk*.

**Supplementary Figure 3** *Igll1*^-/-^ mice maintain both functionally distinct PC1^lo^ and PC1^hi^ subsets of B-1a cells. Representative flow cytometry plots of peritoneal cavity cells from wild-type and *Igll1*^-/-^ mice. B-2 and B-1 cell gates are displayed as a percentage of CD19^+^ cells (left). B-1a and B-1b cell gates are displayed as a percentage of the B-1 cells (middle). PC1lo and PC1 high B-1a cells are displayed as a percentage of B-1a cells (right). A PC1 FMO (fluorescence minus one) was used to help designate the gating.

